# mito-overview: a mitochondrial report software

**DOI:** 10.64898/2026.10.27.762311

**Authors:** Elisson Lopes, Khushi Ahuja, Xiaowu Gai

## Abstract

Interpreting mitochondrial DNA variation requires relating allele representation and genomic annotation to the sequencing evidence available for each sample. We developed mito-overview to integrate mitochondrial coverage, candidate allele fractions, genomic annotations and long read co-occurrence in mitochondrial genomic reports. Analysis of a subset of public GM12878 long read data from Vandiver and colleagues’ Cas9 enrichment study of full-length mitochondrial genomes (SRR18110025) identified 16 candidate variants. Among these candidates, co-occurrence analysis showed that m.8860A>G and m.14831G>A occurred together in all 366 jointly callable query names, demonstrating how mito-overview extends individual allele measurements to quantify shared read support between mitochondrial sites. Analysis of public GM11906 short read data (GSE142741), pooled from three single-cell ATAC-seq libraries, recovered the previously characterized disease-associated m.8344A>G variant in MT-TK at an alternate allele fraction of 72.1% (740 of 1,027 callable observations). The recovered variant identity and MT-TK assignment agree with Lareau and colleagues’ original study, providing a disease-relevant example of mito-overview’s combined candidate detection and genomic annotation. We also examined ONT data from blood, fibroblast and muscle of one patient using a separate internal workflow. Heteroplasmy was estimated from alternate allele fractions calculated separately for each tissue. The respective analyses yielded 90, 52 and 60 candidates, with 19 candidate identities shared across all three lists. For these shared candidates, the tissue-specific estimates enabled examination of both variant presence and allele representation across the sampled tissues, illustrating the application of mitochondrial genomic reports to heteroplasmy assessment in disease research. By combining candidate identification with allele measurements, genomic annotations and shared read support, mito-overview enables researchers to examine which variants are observed, their representation within a sample and their co-occurrence with other mitochondrial alleles. These complementary measurements support the selection and characterization of candidate variants for further study in mitochondrial disease research.

## Introduction

Pathogenic variants in mitochondrial DNA (mtDNA) contribute to disorders with diverse clinical manifestations. Their interpretation is influenced by heteroplasmy, the coexistence of distinct mtDNA alleles within an individual, and by differences in variant abundance among tissues [1]. Investigating mtDNA variation in mitochondrial disease therefore requires characterizing both the sequence changes present and their proportions in the tissues examined. Genomic location and functional annotation provide further context for investigating the biological significance of these variants.

Sequencing enables the detection of mitochondrial variants and estimation of their allele fractions. In nanopore data, coverage and sequencing errors influence these estimates [2]. Long reads also provide information about the arrangement of variants within mitochondrial molecules and can span large deletions [3,4]. For variants at different positions, reads covering both sites allow their co-occurrence to be examined alongside the allele fraction measured at each position. Interpreting these observations requires access to the supporting reads and counts, together with the genomic annotations associated with each candidate.

Here we present mito-overview, a workflow for generating mitochondrial genomic reports from short and long read sequencing alignments. The reports bring together mitochondrial coverage, candidate allele fractions, genomic annotations and, for long read data, co-occurrence among selected variants. Allele measurements are presented with their supporting counts, allowing users to examine the sequence evidence underlying each candidate and its relationship to other mitochondrial alleles. We illustrate these functions using public sequencing datasets and an application of mito-overview to data from multiple tissues from one patient, with the aim of supporting the investigation of mitochondrial variation and heteroplasmy in disease research.

## Materials and Methods

### Workflow inputs and report composition

mito-overview requires an indexed BAM or CRAM alignment, an indexed reference FASTA and a sample configuration. The configuration identifies the input paths, mitochondrial contig, sample identifier, run name and working directory. The workflow is executed through a command line interface in the provided Conda environment. It validates the inputs and extracts the mitochondrial alignments for analysis (Figure 1A).

**Figure 1.**
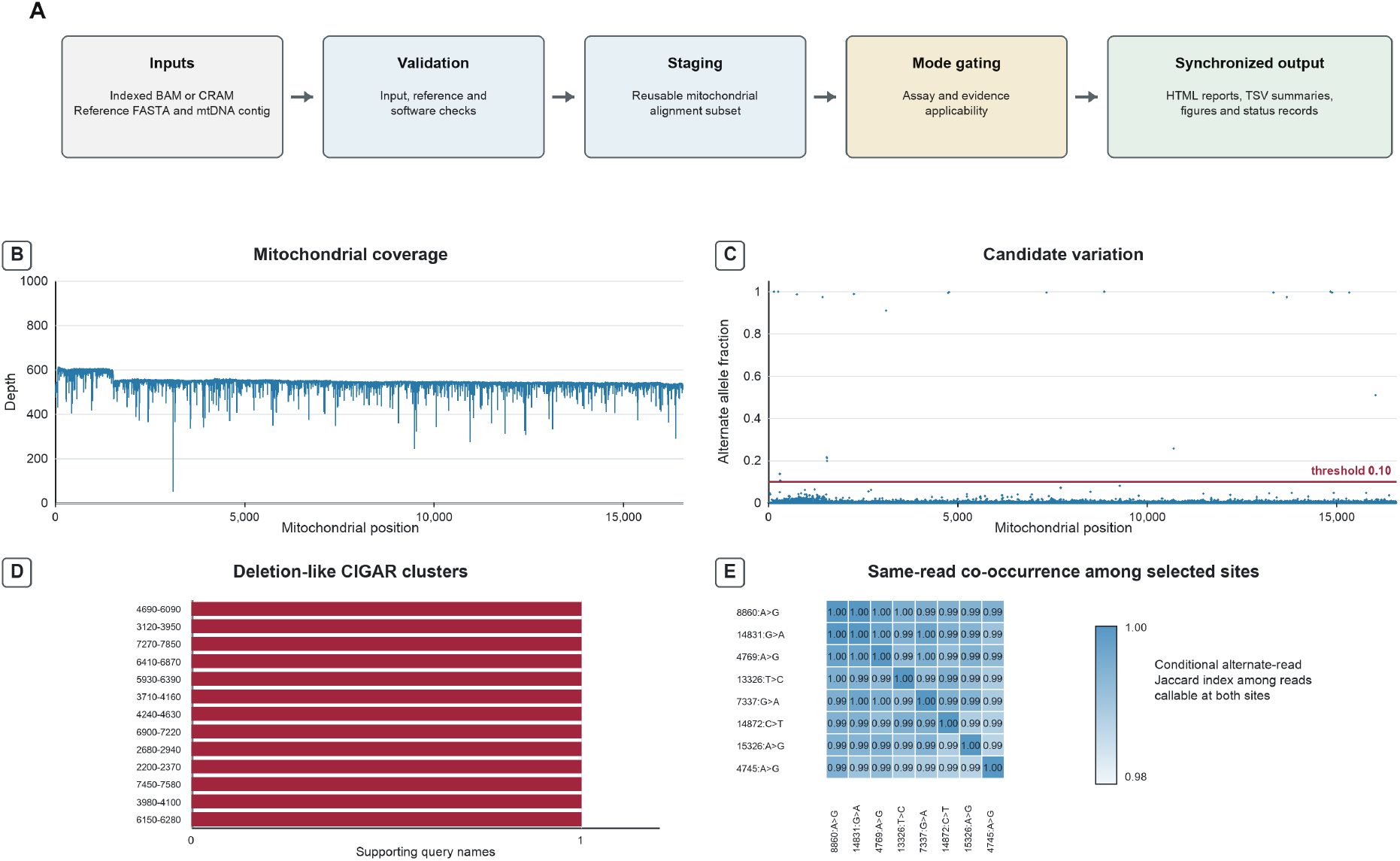
Workflow and GM12878 targeted ONT report. The data panels show mito-overview outputs generated from reads corresponding to 1,000 query names selected from the published GM12878 targeted ONT dataset [4]. (A) Indexed alignment and reference inputs pass validation, mitochondrial staging and assay-mode gating before synchronized report products are assembled. (B) Mitochondrial depth across NC_012920.1. (C) Alternate-allele fractions across the reference, including noncandidate positions: 22 positions were at or above 0.10; 16 met the full candidate criteria, including a canonical reference base and minimum callable depth of 100. The six excluded positions are detailed in Supplementary Table S2. (D) Thirteen binned deletion-like CIGAR patterns from five distinct query names, shown by interval and supporting-name count; these are screening signals, not confirmed deletions. (E) Conditional alternate-support Jaccard among eight selected candidate sites, using query-name sets callable at both sites; the heatmap does not establish phasing.

Read mode and assay type are configured independently, allowing short or long read inputs to be processed under whole genome or targeted mitochondrial profiles. Users can adjust the minimum callable depth and alternate allele fraction for candidate reporting, together with base quality, mapping quality and mean read quality filters. Gene annotations, ClinVar records and ONT modification tracks can be supplied as additional inputs. These resources are incorporated when they meet the requirements of the selected analysis and reference.

Both read modes provide coverage, candidate and compatible annotation summaries. Long read mode also evaluates co-occurrence among selected candidates and screens alignments for deletion-like patterns.

Additional analyses depend on the assay and available reference information. Targeted mitochondrial profiles exclude the experimental mitochondrial to nuclear depth ratio and the Phy-Mer haplogroup interface. The reference is assessed separately to determine whether alignment ambiguity involving nuclear mitochondrial sequences (NUMTs) can be assigned a caution category.

The outputs are organized into HTML report pages, numerical TSV tables and available figures. Candidate summaries include the alternate counts and total callable observations used to calculate each allele fraction. Co-occurrence summaries provide the counts available at both selected positions and the support for their alternate alleles. Each analysis has a status indicating its outcome and whether its input, assay and evidence requirements were met. On completion, the report products are collected with the configuration and records of the analysis. The Supplementary Reproducibility Note provides the exact invocation, full profile rules, reference checks and output inventory. Public long and short read datasets were analyzed with mito-overview v0.3.1. A patient application used preserved internal analysis and report scripts for blood, fibroblast and muscle inputs, with the implementation and settings described below.

Optional capabilities include an experimental mitochondrial to nuclear depth ratio, modification summaries from supplied ONT bedMethyl tracks, a Phy-Mer haplogroup interface and external candidate annotation through mvTool. The depth ratio is dimensionless rather than an absolute mitochondrial copy count, and modification summaries remain exploratory [5,6]. These capabilities were not illustrated in the public Results. Their calculations, applicability rules and interface checks are described in the Supplementary Reproducibility Note.

### Mitochondrial coverage and alignment assessment

Mitochondrial QC coverage is calculated at each reference position by summing A, C, G and T observations obtained with pysam count_coverage at a base quality threshold of zero and its default read callback. Mean and median depths summarize the resulting profile across the mitochondrial reference, including positions with zero coverage. This QC profile describes the coverage background for the sample; the filtered callable depth used for allele fraction calculation is defined separately below (Figures 1B and 2A).

For long read inputs, seven thresholded alignment conditions are combined into a heuristic NUMT caution score: low and very low mapping quality, short aligned spans, extensive soft clipping, supplementary alignment tags, supplementary alignment records, and scarcity of nearly complete primary alignments. Each satisfied condition contributes one point. Scores of 0-1, 2-3 and 4-7 correspond to low, moderate and high caution, respectively. Category assignment requires a supported whole genome reference profile and complete, internally consistent alignment metrics. Computable metrics remain available when a category cannot be assigned. The score describes alignment properties at the sample level; the component thresholds, alignment-record denominators and reference requirements are specified in the Supplementary Reproducibility Note.

### Allele fraction calculation and candidate selection

mito-overview calculates allele fractions from A, C, G and T observations that pass configurable alignment and base filters. Default minimum base quality, mapping quality and mean read quality are 13, 20 and 10, respectively. Mean read quality is the arithmetic mean of recorded base quality scores. Records matching SAM exclusion mask 3844 are removed, as are observations corresponding to deletions, reference skips, missing bases or noncanonical bases. With overlap suppression enabled by default, at most one passing observation per query name is retained at each position. The precise overlap selection, quality handling and depth-cap rules are specified in the Supplementary Reproducibility Note.

At mitochondrial position i, the retained counts A_i_, C_i_, G_i_ and T_i_ define callable depth d_i_ (Figure 2C,D). The most abundant single nonreference base is selected as the alternate allele, and its count a_i_ is divided by callable depth to obtain the alternate allele fraction h_i_:

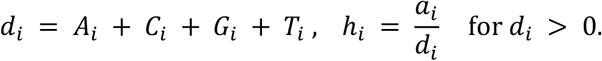

**Figure 2.**
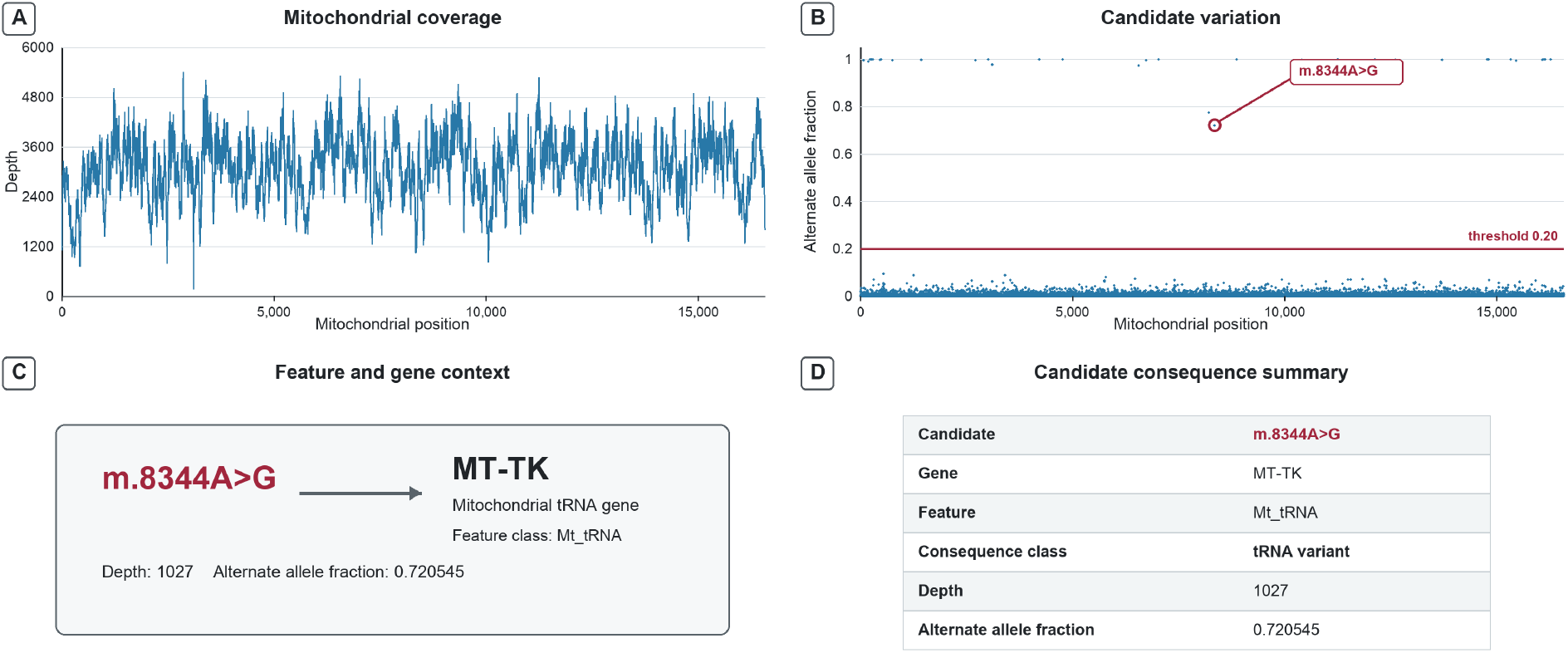
Pooled public GM11906 short-read report. The three GM11906 single-cell ATAC-seq libraries are from the published dataset [7]; their reads were pooled for this report. (A) Mitochondrial depth across NC_012920.1. (B) Alternate-allele fractions include noncandidate positions; 33 sites met both minimum callable depth 10 and fraction threshold 0.20. The previously characterized m.8344A>G site [7,8,9] is highlighted, with 740 alternate observations among 1,027 callable observations in the pooled input. (C) Feature and gene context places that site in MT-TK with feature class Mt_tRNA. (D) The candidate consequence summary labels it tRNA_variant.

Candidate assessment requires an A, C, G or T reference base and positive callable depth meeting the configured minimum. A site is reported when the selected alternate allele has a positive count and both reporting thresholds are satisfied:

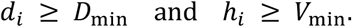

The package defaults are D_min_=100 callable observations and V_min_=0.02; the public examples use the settings specified under Application datasets and analysis settings. Positions with zero callable depth have an undefined fraction. When no position is eligible, the candidate screen is unevaluable. Otherwise, a result with no reported candidates applies to the eligible positions under the configured thresholds.

### Read co-occurrence and deletion screening

Candidate co-occurrence is evaluated using the alignment and base filters described above. The module selects up to eight candidate sites, ranked first by decreasing alternate fraction, then by decreasing callable depth, with increasing coordinate resolving remaining ties. Under default overlap suppression, each site pair is evaluated among query names with passing observations at both positions, requiring at least 25 such names. Within this shared set, n_alt1_ and n_alt2_ count names supporting each selected alternate, and n_both_ counts names supporting both. The conditional alternate-support Jaccard index is

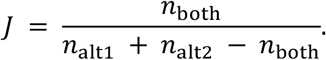

Thus, among jointly callable query names supporting at least one selected alternate, J measures the proportion supporting both. A zero alternate-support union leaves the index undefined. Fewer than two valid selected sites, no pair meeting the shared-name requirement or no pair with a defined index leaves the analysis unevaluable.

The deletion screen identifies large CIGAR deletion operations in mapped primary and supplementary mitochondrial alignments, excluding unmapped and secondary records. This screen uses a separate alignment selection from the allele fraction and co-occurrence analyses. Events meeting the configured minimum length, defaulting to 100 bp, are grouped by their start and end coordinates into fixed 10 bp bins. Each bin pair reports the number of distinct supporting query names.

For each bin pair, the support fraction is calculated by dividing the number of supporting query names that also have a retained primary mitochondrial alignment by the total number of distinct query names with such a primary alignment. An event on a supplementary segment contributes to this fraction when the same query name also has a retained primary mitochondrial alignment. With no retained primary names, the fraction is undefined.

### Genomic annotation and gene summaries

mito-overview annotates candidate variants by linking their callable depths and alternate allele fractions to mitochondrial genomic features. In human reports, candidate positions are intersected with gene and coding sequence intervals from the configured mitochondrial GTF, retaining overlapping feature assignments. These assignments distinguish protein coding genes, tRNAs and rRNAs, while additional positional annotations identify control region and intergenic sites. Reference requirements for control region annotation are described in the Supplementary Reproducibility Note.

For candidates within protein coding genes, the annotated coordinates and strand define the reference codon containing the substituted base. The selected alternate base is introduced into that codon, and the reference and alternate sequences are translated using the vertebrate mitochondrial genetic code. Comparison of the translated codons yields synonymous or missense assignments, with separate handling of changes involving initiation or termination codons. Detailed protein changes are calculated for complete codons within a single contiguous coding sequence annotated with phase zero.

When a ClinVar VCF is supplied, candidate variants are matched by contig, position, reference and alternate allele to retrieve recorded clinical significance and disease annotations. Clinical variant classification remains a separate interpretive step [1]. At the gene level, summaries report the number of candidate positions, median callable depth at those positions, and mean and maximum alternate allele fractions. Counts of sites selected for co-occurrence analysis and overlaps with deletion events relate the sequence and structural screening outputs to the same genomic features.

### Application datasets and analysis settings

The public examples used mito-overview v0.3.1. The long read example used targeted ONT GM12878 data from PRJNA809571/SRR18110025 [4] to examine candidate eligibility, support shared between mitochondrial sites and deletion-like alignment patterns. We selected 1,000 query names deterministically by seeded SHA-256 ranking and aligned their reads to the mitochondrial reference NC_012920.1 with minimap2 in map-ont mode. The report used the long read, targeted mitochondrial profile, with minimum callable depth 100 and minimum alternate allele fraction 0.10 for candidate reporting.

The short read example used the previously characterized GM11906 m.8344A>G marker [7,8,9] to examine the connection between callable support, alternate fraction and genomic annotation. We pooled paired reads from three GM11906 single-cell ATAC-seq libraries [7] (SRR10804585, SRR10804590 and SRR10804657), concatenating their read-1 and read-2 files separately, and aligned the pool to NC_012920.1 with BWA-MEM. For this mitochondrial alignment, we used the short read, targeted mitochondrial report profile, with minimum callable depth 10 and minimum alternate allele fraction 0.20. Candidate fractions describe passing callable observations in the pooled input, not per-cell estimates.

The patient application examined mitochondrial candidate identity and allele representation in existing barcode-resolved ONT alignments from blood, fibroblast and muscle from one individual. Mitochondrial alignments were extracted from each indexed BAM with samtools and analyzed using the GRCh38 primary-assembly reference and the preserved analysis and report scripts documented in the Supplementary Reproducibility Note. For this application, candidate sites were calculated from A, C, G and T observations counted with pysam at a base quality threshold of zero and without read callback filtering. Candidate depth was the sum of these counts, and the alternate allele fraction was the single highest nonreference count divided by that depth. Candidates required depth of at least 100 and alternate allele fraction of at least 0.02. These observation settings differ from the released workflow defaults described above.

Candidate lists were matched across tissues by mitochondrial position, reference allele and selected alternate allele. For shared candidate identities, allele fractions and their associated depths were retained separately for each tissue. Entries absent from a tissue’s candidate list were left unavailable in the candidate-derived comparison rather than assigned a zero allele fraction. Tissue QC coverage was calculated separately at the same base quality threshold using the default read callback. Candidate annotation summaries counted rows with coding-consequence or feature/location labels, and each tissue’s coverage profile was paired with its annotation summary on independently scaled axes (Figure 3). A separate deletion-like screen identified CIGAR D segments of at least 100 bp. The Supplementary Reproducibility Note records the case-specific implementation, preparation and annotation procedures.

**Figure 3.**
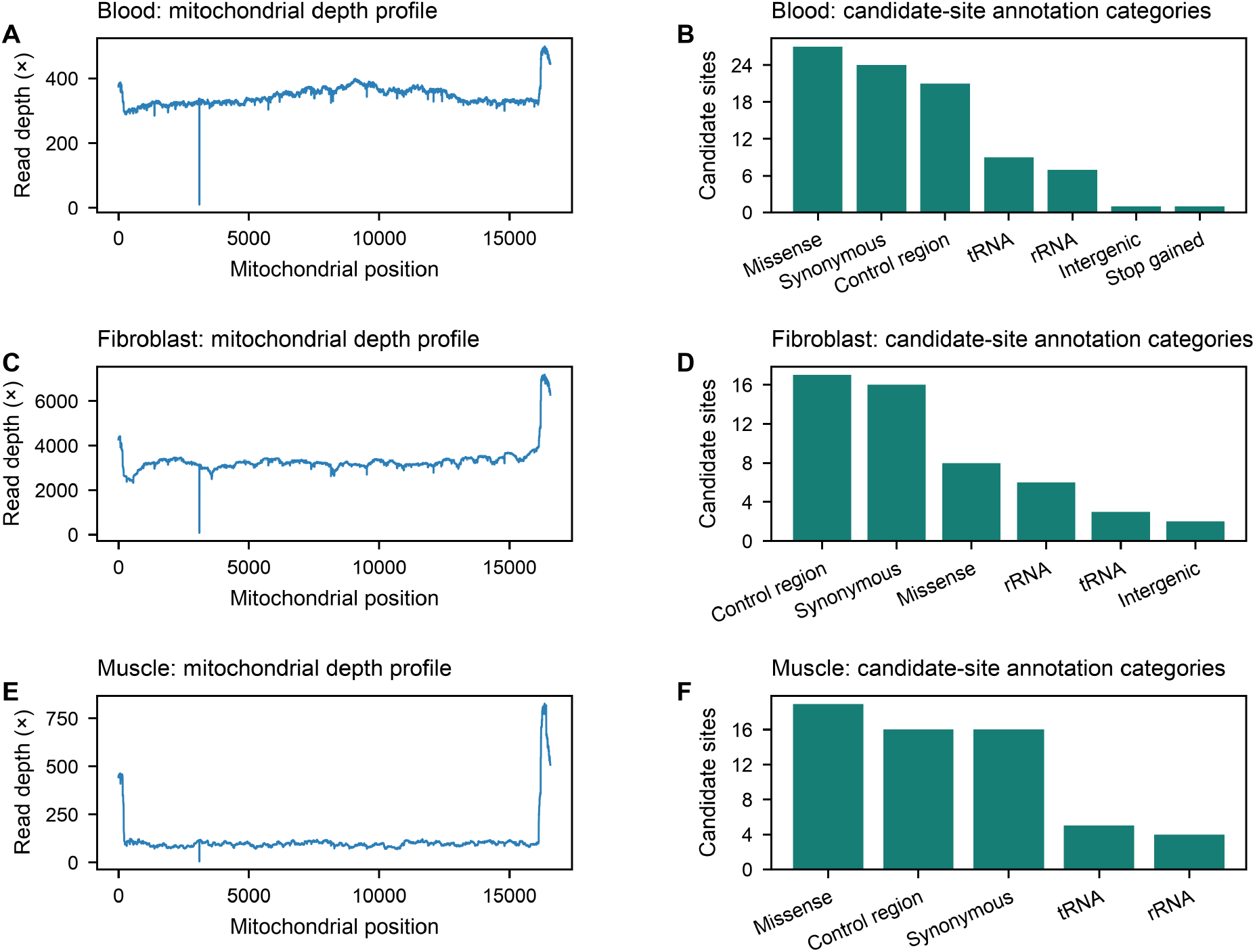
Coverage and candidate annotations across patient tissues. Reports from barcode-linked blood, fibroblast and muscle inputs from one individual provide coverage and annotation context for the allele fractions in Table 1. The patient application used the preserved mito-overview implementation described in Methods. Panels A, C and E show QC mitochondrial depth across position for blood, fibroblast and muscle, respectively; panels B, D and F show counts of annotated candidate rows under mixed coding-consequence and feature/location labels. Candidate sites meet reporting thresholds of a minimum A/C/G/T observation depth of 100 and a minimum alternate allele fraction of 0.02. The QC and candidate counters use separate read callback settings, as described in Methods. Panel y-axes are scaled independently.

### Ethics and Data Use

The public examples used previously deposited sequencing data and public cell-line metadata. The patient application used existing ONT sequencing data from blood, fibroblast and muscle of one individual. Only summary results from these analyses are presented.

**Table 1.** Tissue heteroplasmy estimates for two shared mitochondrial candidates.

| Candidate | Tissue | Alternate observations | Candidate depth | Alternate allele fraction (%) |
| --- | --- | --- | --- | --- |
| m.16265A>G | Blood | 141 | 486 | 29.0 |
| m.16265A>G | Fibroblast | 2,569 | 7,076 | 36.3 |
| m.16265A>G | Muscle | 219 | 788 | 27.8 |
| m.73A>G | Blood | 350 | 358 | 97.8 |
| m.73A>G | Fibroblast | 3,882 | 3,912 | 99.2 |
| m.73A>G | Muscle | 423 | 425 | 99.5 |

Values were obtained using the patient application of mito-overview described in Methods. Candidate depth is the total A, C, G and T observations at each site. Alternate allele fractions are alternate counts divided by candidate depth, expressed as percentages rounded to one decimal place. All tissues are from one individual.

### Software evaluation and reproducibility

Evaluation of mito-overview v0.3.1 addressed expected software behavior, filter dependence and reproducibility. Recorded unit and known-answer tests exercised filtered allele counting, candidate eligibility and threshold behavior, and denominator and undefined-value handling. Four synthetic or standalone smoke-test cases exercised workflow execution and reporting, including profile-dependent applicability and missing-input states. For each public input, two reports with default observation filters tested output consistency, while one report with lenient filters and one with strict filters examined filter dependence at fixed candidate thresholds, giving eight report executions in total. Minimum base quality, mapping quality and mean read quality were 13, 20 and 10 for the default profile; 0, 0 and 0 for the lenient profile; and 20, 30 and 15 for the strict profile. Numerical summaries and report states were compared with frozen expected values. Repeated default reports, using the same derived alignment, were compared through normalized TSV tables, HTML structure and decoded PNG pixels.

To assess reproducibility after independent installation and alignment reconstruction, two independently executed workflows each started from a fresh installation of the same release and byte-identical copies of the seven public FASTQ files, regenerated the two example alignments, and repeated the report matrix against the same frozen expected values. These checks assessed reproducibility for the public inputs under the tested configurations. The patient tissue application was not part of this release evaluation. Execution details and supporting records are described in the Supplementary Reproducibility Note.

## Results

### Characterization of mitochondrial variants in public sequencing data

mito-overview identified 16 mitochondrial candidate sites in the GM12878 targeted ONT subset comprising 1,000 query names [4] (Figure 1C). Among these candidates, m.8860A>G and m.14831G>A were annotated as missense variants in MT-ATP6 (p.T112A) and MT-CYB (p.A29T), respectively. Co-occurrence analysis linked these annotated candidates through the reads supporting both alternate alleles, extending their characterization beyond separate sitewise measurements.

Co-occurrence was evaluable for all 28 pairs formed by eight selected candidate sites (Figure 1E). For the report-selected strongest pair, m.8860A>G and m.14831G>A, the individual alternate fractions were both 1.0, based on 474 and 433 callable observations, respectively. Within the shared set of 366 query names with passing observations at both positions, all supported both alternate alleles, yielding a conditional alternate-support Jaccard value of 1.0.

Candidate assessment was possible at 15,723 of the 16,569 reference positions; the separate QC coverage profile had a mean depth of 545.484 and a median of 544 (Figure 1B). Of 22 positions reaching an alternate fraction of 0.10, 16 met the complete candidate criteria, including a canonical reference base and at least 100 callable observations. The remaining six positions had fewer than 100 callable observations. At position 152, for example, all 96 callable observations supported the alternate allele, but the site did not meet the minimum depth. Position 3107 additionally had a noncanonical reference base, N. The six excluded positions and their unmet criteria are listed in Supplementary Table S2.

The alignment-based structural screen provided a distinct set of observations from the same input. It reported 13 deletion-like CIGAR interval bins, each with one supporting query name; five distinct query names accounted for support across all 13 bins (Figure 1D). The report also retained mapping quality, alignment span and clipping metrics. Because the alignment reference contained only NC_012920.1, these metrics were available without a categorical NUMT caution assignment.

GM11906 provided a complementary short read application in which a disease-associated mitochondrial allele had been characterized previously [7,8,9]. Among 33 candidates meeting minimum callable depth 10 and minimum alternate fraction 0.20, mito-overview identified m.8344A>G with 740 alternate observations among 1,027 callable observations, corresponding to an alternate fraction of 0.720545, or 72.1% (Figure 2B). The same candidate was assigned to MT-TK in the feature and consequence outputs, with feature class Mt_tRNA and consequence tRNA_variant (Figure 2C,D). Its identity agrees with the marker studied by Lareau and colleagues [7], and its gene assignment corresponds to the established mitochondrial tRNA-Lys context [8,9].

The GM11906 measurements describe passing observations pooled from three single-cell ATAC-seq libraries, rather than estimates for individual cells. The accompanying QC coverage profile had a mean mitochondrial depth of 3,098.6 and a median of 3,144 (Figure 2A). The selected short read, targeted mitochondrial profile retained the candidate and annotation analyses and marked co-occurrence, deletion screening, NUMT quality control and the mitochondrial to nuclear depth ratio as not applicable.

### Mitochondrial allele representation across patient tissues

The patient application used the preserved implementation described in Methods to examine matched mitochondrial candidates across blood, fibroblast and muscle from one individual. Under common candidate criteria of depth at least 100 and alternate fraction at least 0.02, the three reports contained 90, 52 and 60 candidate sites, respectively. These lists were obtained from inputs with mean mitochondrial QC depths of 346.413, 3,330.269 and 114.574 in the same tissue order (Figure 3A,C,E). Matching the candidate tables by position, reference and alternate allele identified 134 distinct candidate identities across the lists, of which 19 were reported in all three tissues.

The measurements at shared sites provided further information beyond their inclusion in all three candidate lists (Table 1). The m.16265A>G candidate, annotated to the mitochondrial control region in each report, had an alternate allele fraction of 29.0% in blood, supported by 141 alternate observations among 486; 36.3% in fibroblast, supported by 2,569 among 7,076; and 27.8% in muscle, supported by 219 among 788. Another shared candidate, m.73A>G, had alternate fractions close to 1 in all three tissues: 97.8%, 99.2% and 99.5%, respectively.

Candidate annotations added genomic context to the measurements in each tissue. Missense and synonymous consequences, together with control region, tRNA and rRNA labels, were represented in all three reports (Figure 3B,D,F). Figure 3 pairs these annotation summaries with the corresponding QC coverage profiles on independently scaled axes. The candidate membership and deletion-screen summaries are detailed in the Supplementary Reproducibility Note.

### Reproducibility of candidate measurements and annotations

The public-data evaluation distinguished consistency of repeated reports from agreement with expected outputs under different observation filters. Across eight executions of the two public inputs, numerical summaries, output inventories and assay-dependent report states matched the expected results for their respective configurations. In repeated default analyses of the pooled GM11906 input, the m.8344A>G candidate row, its feature and consequence annotations, and accompanying report states agreed. All 36 recorded release validation cases passed, including four synthetic or standalone smoke-test cases, and the original release unit-test suite passed 1,256 tests.

Reproducibility after rebuilding the analysis inputs was assessed in two independently installed workflows that regenerated the alignments from the same public FASTQ files. Each passed all 17 public-matrix cases and all 366 assertions against the frozen expected values. Within each workflow, repeated default reports agreed in normalized numerical tables and module states, HTML structure and decoded PNG pixels. Each independent installation’s final unit-test suite recorded 1,254 passed tests, two skips and no failures. The Supplementary Reproducibility Note distinguishes these outcomes from the original release evaluation and documents the skipped tests.

## Discussion

mito-overview supports mitochondrial disease research by combining candidate detection and heteroplasmy estimation with genomic annotation and examination of supporting reads. These functions address complementary questions in the investigation of mtDNA disorders. For example, studying the m.8344A>G allele associated with myoclonic epilepsy with ragged red fibers (MERRF) requires characterizing its abundance as well as its identity [8]. Investigations of disorders involving large mtDNA deletions, including some forms of chronic progressive external ophthalmoplegia, additionally require structural evidence [3]. Within this research setting, mito-overview connects quantitative assessment of sequence variants with inspection of their genomic context and, for long reads, their co-occurrence and deletion-like alignment patterns.

The public applications demonstrate how this approach accommodates mitochondrial data from different experimental designs. In targeted ONT data from GM12878, mito-overview identified 16 candidates and linked variants in MT-ATP6 and MT-CYB through their shared read support, extending individual allele measurements to the examination of co-occurrence. In pooled GM11906 single-cell ATAC-seq data, it recovered m.8344A>G at an alternate fraction of 72.1% and assigned it to MT-TK. The recovered identity and gene assignment agree with the marker characterized in the original study [7-9]. This application demonstrates the use of mitochondrial reads from an assay designed for chromatin accessibility to quantify a previously characterized disease allele. The workflow retained candidate quantification and annotation for both inputs while applying analyses appropriate to each read mode. Reproduction of the public candidate measurements and annotations after independent installation and alignment reconstruction further supports the computational consistency of these analyses.

Application of mito-overview to ONT data from blood, fibroblast and muscle of one patient extended this analysis to heteroplasmy across sampled tissues. Among the 19 candidate identities shared across all three reports, m.16265A>G had estimated alternate fractions of 29.0%, 36.3% and 27.8%, respectively, whereas m.73A>G was represented at 97.8%-99.5%. The measurements therefore characterized both intermediate heteroplasmy and nearly complete alternate representation within the same patient, with counts and local depths available for examining each estimate. This is relevant to mitochondrial disease investigation because heteroplasmy measured in one tissue may not represent its level in another, including a tissue affected by disease [1].

Interpretation of low abundance candidates remains an area for further calibration in mito-overview. For the public analyses, allele fractions are calculated from filtered observations, and the reporting thresholds define candidate selection rather than an established limit of detection. Evaluation with independently characterized mixtures could support uncertainty estimates and refinement of these thresholds across sequencing conditions. The deletion and NUMT summaries provide screening evidence from the supplied alignments, with detailed event characterization requiring complementary approaches. Optional plugins could incorporate splice-aware deletion evidence from NanoDel [10] and, for whole genome long read data, nuclear insertion calls from ANOMALY [11]. Testing these integrations against characterized events would establish their contribution to structural interpretation.

The central contribution of mito-overview is to connect mitochondrial sequence variation with its quantitative and genomic context in the biological samples under investigation. By linking candidate identity and abundance to supporting sequence evidence, the workflow provides a basis for selecting variants for molecular confirmation and subsequent investigation of their relationship to tissue phenotypes and mitochondrial function.

## Data and code availability

mito-overview v0.3.1, which retains scientific protocol v0.3.0, is available under the MIT license at the tagged GitHub release (https://github.com/elissonnog/mito-overview/releases/tag/v0.3.1). The release provides a validation packet (https://github.com/elissonnog/mito-overview/releases/download/v0.3.1/mito-overview-v0.3.1-validation.zip) and validation report (https://github.com/elissonnog/mito-overview/releases/download/v0.3.1/MitoOverview_v0.3.1_release_validation_report.pdf). Public sequencing inputs used here are GM12878 run SRR18110025 and GM11906 runs SRR10804585, SRR10804590 and SRR10804657. The records supporting the later independent executions are retained locally, are separate from the release validation packet and have no public archive specified here. The Supplementary Reproducibility Note identifies these records and distinguishes report-native repeatability from the manuscript figure composites. The underlying sequencing data for the separate one-case tissue application are controlled internal material and are not publicly available.

## Notes

https://github.com/elissonnog/mito-overview

### Competing Interest Statement

The authors have declared no competing interest.

## References

1. McCormick EM, Lott MT, Dulik MC, et al. Specifications of the ACMG/AMP standards and guidelines for mitochondrial DNA variant interpretation. Hum Mutat. 2020;41(12):2028–2057. doi: 10.1002/humu.24107. https://pubmed.ncbi.nlm.nih.gov/32906214/

2. Slapnik B, Sket R, Crepinsek K, et al. The quality and detection limits of mitochondrial heteroplasmy by long read nanopore sequencing. Sci Rep. 2024;14:26778. doi: 10.1038/s41598-024-78270-0. https://www.nature.com/articles/s41598-024-78270-0

3. Frascarelli C, Zanetti N, Nasca A, et al. Nanopore long-read next-generation sequencing for detection of mitochondrial DNA large-scale deletions. Front Genet. 2023;14:1089956. doi: 10.3389/fgene.2023.1089956. https://pubmed.ncbi.nlm.nih.gov/37456669/

4. Vandiver AR, Pielstick B, Gilpatrick T, et al. Long read mitochondrial genome sequencing using Cas9-guided adaptor ligation. Mitochondrion. 2022;65:176–183. doi: 10.1016/j.mito.2022.06.003. https://pubmed.ncbi.nlm.nih.gov/35787470/

5. Bicci I, Calabrese C, Golder ZJ, Gomez-Duran A, Chinnery PF. Single-molecule mitochondrial DNA sequencing shows no evidence of CpG methylation in human cells and tissues. Nucleic Acids Res. 2021;49(22):12757–12768. doi: 10.1093/nar/gkab1179. https://pubmed.ncbi.nlm.nih.gov/34850165/

6. Guitton R, Nido GS, Tzoulis C. No evidence of extensive non-CpG methylation in mtDNA. Nucleic Acids Res. 2022;50(16):9190–9194. doi: 10.1093/nar/gkac701. https://pubmed.ncbi.nlm.nih.gov/35979955/

7. Lareau CA, Ludwig LS, Muus C, et al. Massively parallel single-cell mitochondrial DNA genotyping and chromatin profiling. Nat Biotechnol. 2021;39:451–461. doi: 10.1038/s41587-020-0645-6. https://www.nature.com/articles/s41587-020-0645-6

8. Shoffner JM, Lott MT, Lezza AMS, et al. Myoclonic epilepsy and ragged-red fiber disease (MERRF) is associated with a mitochondrial DNA tRNA(Lys) mutation. Cell. 1990;61(6):931–937. doi: 10.1016/0092-8674(90)90059-N. https://pubmed.ncbi.nlm.nih.gov/2112427/

9. Coriell Institute for Medical Research. GM11906 - B-Lymphocyte sample record. Accessed 2026 Oct 7. https://www.coriell.org/0/Sections/Search/Sample_Detail.aspx?Ref=GM11906

10. Fearn C, Poulton J, Fratter C, Oliva C, Griguer C, Baldock RA, et al. “NanoDel”: identification of large-scale mitochondrial DNA deletions using long-read sequencing. Bioinformatics. 2026;42(10):btag684. doi: 10.1093/bioinformatics/btag684.

11. Mahar NS, Singh R, Gupta I, Ramdas S. ANOMALY: a Snakemake pipeline for identifying NuMTs from long-read sequencing data. NAR Genom Bioinform. 2026;8(1):lqag014. doi: 10.1093/nargab/lqag014.

